# Accurate Reconstruction of Circular RNAs from Complex Rolling Circular Long Reads with CircPlex

**DOI:** 10.1101/2025.11.21.689841

**Authors:** Tasfia Zahin, Irtesam Mahmud Khan, Mingfu Shao

## Abstract

Rolling circle amplification (RCA) coupled with long-read sequencing has emerged as a powerful strategy for detecting full-length circular RNAs (circRNAs). Such protocols produce long reads that are normally composed of several tandemly repeated copies of the original circRNA. The circRNA sequence can be reconstructed through detecting the repeating unit of the long reads, which are aligned to the genome to validate and to identify back-splice junctions (BSJs). We revealed a previously unrecognized phenomenon: a substantial fraction of long reads contain complex repeat patterns in which the repeating unit consists of a sequence combined with its partial reverse complement. In these cases, only the original sequence corresponds to the true circRNA, while the concatenated pattern may produce false circRNAs and misidentify correct circRNAs. We present a new approach CircPlex that extracts the authentic circRNA sequence from these complex repeat units, overcoming the limitations of standard repeat-based consensus prediction. Comparison with isoCirc annotations and circRNA database demonstrates that a significant number of BSJs and full-length sequences, previously ignored, can be recovered. Our results suggest that leveraging partially repetitive reads from RCA-based sequencing can substantially increase circRNA detection sensitivity and uncover novel isoforms, providing a more comprehensive view of the circular transcriptome.

## 1 Introduction

Circular RNAs (circRNAs) are a class of covalently closed RNA molecules that form a continuous loop structure. They are generated through a non-canonical splicing process known as *back-splicing*, in which the 3’ end of a downstream exon is joined to the 5’ end of an upstream exon via a back-splicing junction (BSJ), resulting in a stable circular RNA structure. CircRNAs have gained increasing attention for their diverse biological roles [22, 9] and clinical potential [7, 30]. They play important roles in regulating gene expression at multiple levels, ranging from modulating transcription and splicing [45, 16, 3, 44], acting as sponges for multiple microRNAs (miRNAs) by harboring numerous binding sites [27, 37, 4], to influencing mRNA function [32, 1, 33]. CircRNAs are highly resistant to exonuclease degradation and consequently more stable than their linear counterparts. Such properties make them suitable for a wide variety of applications, including functioning as noncoding aptamers, modulating innate immune responses, and serving as antisense RNAs [12, 23, 31]. Many circRNAs show distinct tissue and developmental stage-specific expression patterns, indicating finely tuned regulatory control. Disease-relevant circRNAs are emerging as promising therapeutic targets [15, 36, 30], while their stable and specific expression patterns also make them valuable as diagnostic biomarkers for a range of conditions, including cancers [18, 8, 26], autoimmune diseases, and neurodegenerative disorders [48, 23, 11].

The rapidly growing biological research and therapeutic applications involving circRNAs rely fundamentally on accurately determining their full-length sequences. This is primarily accomplished by RNA sequencing (RNA-seq) coupled with computational approaches that reconstruct full-length circRNAs from the resulting RNA-seq reads. The reconstruction remains an exceedingly difficult task, due to the low expression levels of circRNAs, their extensive overlap with cognate linear transcripts, and the inherent limitations of current sequencing protocols. These challenges highlight the necessity for new computational methods capable of reliably reconstructing full-length circRNAs with high accuracy.

Numerous computational methods have been developed to detect circRNAs from short-read RNA-seq data, leveraging its low error rate and high sequencing depth to identify back-splicing junctions (BSJs). Several of them like CYCLeR [35], psirc [40], CircAST [38], CIRCexplorer2 [43] depend on annotation and does not work well for non-model species. Other tools including CIRI3 [46], CIRI2 [13], Circall [29], CircMiner [2], CircMarker [21], CIRCexplorer3 [25], and Sailfish-cir [20] are effective for BSJ detection and quantification, but they often fall short in reconstructing full-length circRNA sequences. Tools like CIRI-full [47] and TERRACE [41] can assemble full-length circRNAs from short reads; however, their performance remains suboptimal, particularly in reconstructing low-abundance or novel circRNAs. To address these limitations, long-reads sequencing technologies have recently been adopted for circRNA profiling. Among them, isoCirc [39], CIRI-long [42] and circFL-seq [24] use rolling circular techniques, such as rolling circular amplification (RCA) and rolling circular reverse transcription (RCRT), that produce (long) molecules composed of several tandemly repeated copies of the original circRNA template in the library preparation. Followed by long-reads sequencing such as Oxford Nanopore or PacBio [10, 6], these methods generate the so-called rolling circular long reads that contain multiple copies of the captured circRNA but with errors (see an example in Fig. 1a). Reconstructing full-length circRNAs from this type of data amounts to deriving the repeating unit from the error-prone reads. Several methods have been developed for this task, including mTR[28], TRF [5], and EquiRep [34] that we developed.

**Figure 1.**
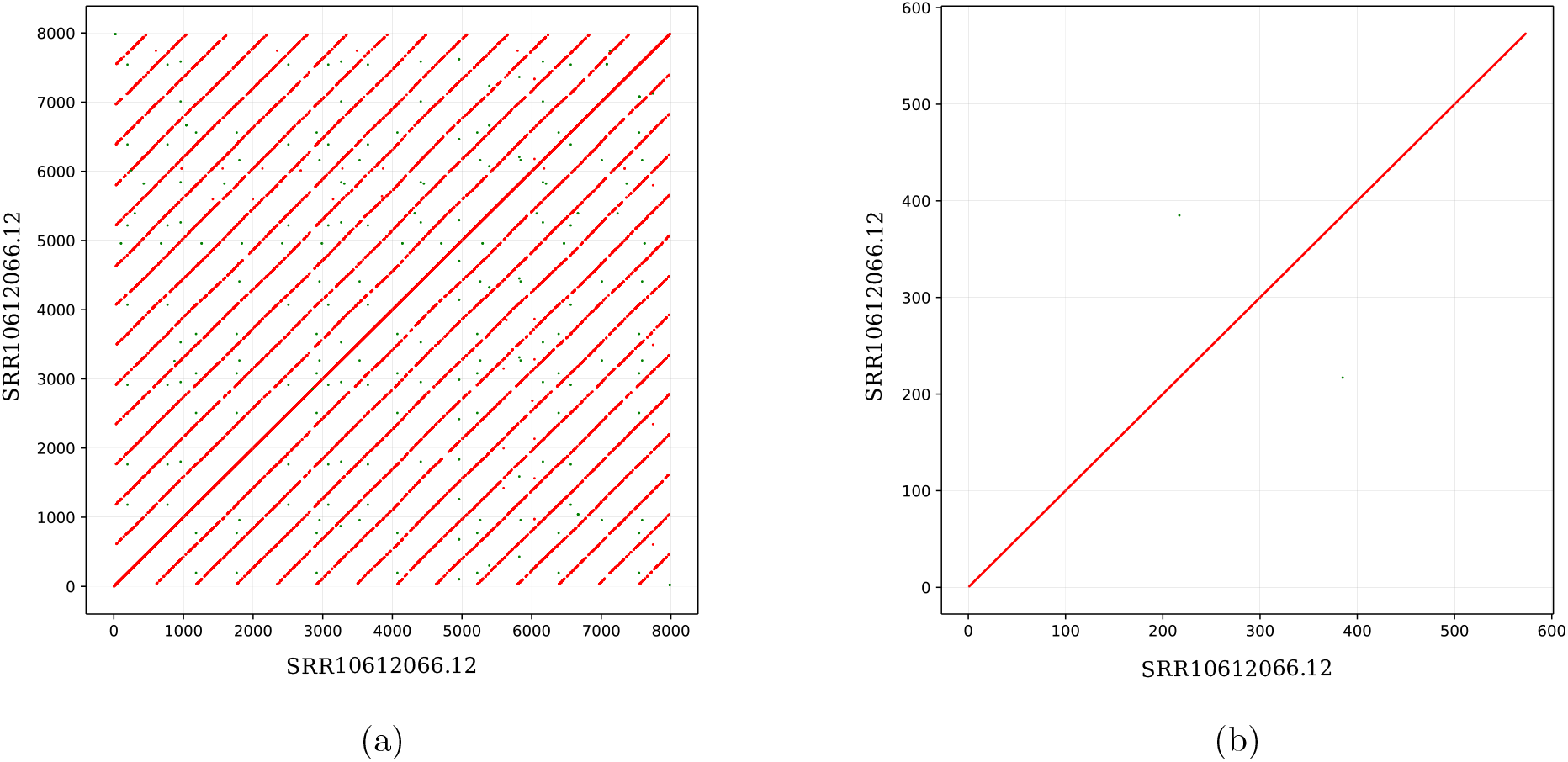
Dotplot of a regular read from the RCA protocol and its unit predicted by EquiRep. Red dots are shared kmers (length 10) between the sequence and itself; green dots are shared kmers (length 10) between the sequence and its reverse complement. (a) Dotplot of read SRR10612066.12; the dotted red lines show the multiple repeating copies of a potential circRNA sequence. (b) Dotplot of the repeating unit predicted by EquiRep from SRR10612066.12.

As we investigate circRNAs more deeply, we discovered that many rolling circular long reads exhibit a complex repeating patterns, where the repeating unit contains the circRNA followed by its partial reverse complement. Consequently, the constructed repeating unit from these complex long reads, which were treated as the circRNA sequences, are in fact incorrect. In this work, we describe this previously unrecognized phenomenon and propose a new method CircPlex that accurately reconstructs genuine circRNA sequences from such complex reads. Our approach begins by determining whether a read might be complex using kmer-profiling; for such reads, we extract the repeat unit using existing methods and design a new algorithm to precisely reconstruct the true circRNAs. Specifically, we first perform systematic rearrangements to transform the unit into a standardized configuration, and then apply an algorithm to identify partitions that separate the true circRNA sequence from its partial reverse complement. We validate our predicted sequences by aligning to the reference genome and comparing their BSJs to existing databases. Results confirm our conjecture regarding the pattern in complex reads and demonstrate that the sequences we reconstruct reliably correspond to genuine circRNAs, some of which are novel circRNAs that are overlooked by conventional pipelines. We anticipate that the principles we highlight, together with the assembly framework we introduce, will be broadly adopted in future circRNA research.

## 2 Methods

Given a sequenced long read from the RCA protocol, CircPlex employs a four-step approach to determine if it is a complex read and if yes, reconstructs the true circRNA sequence.

### 2.1 Identifying potentially complex RCA long read

A long read generated using the RCA protocol often exhibits a regular pattern, in which multiple imperfect copies of the circRNA sequence appear in tandem (Fig. 1a shows an example). However, some reads show a distinctive pattern where each repeat consists of the circRNA sequence followed by a partial reverse complement of itself. We refer to these as complex reads (Fig. 2a shows an example). The first step of CircPlex is to determine whether an input read is regular or complex. To achieve this, we employ a kmer-based approach. We use *KMC* [17] to count all non-canonical kmers in the read. Kmers with counts less than 2 are discarded, and the total number of remaining kmers is denoted as *K*. Each kmer is then paired with its reverse complement (if present in the count file), and we sum the minimum count between each pair, yielding a value *R*. If the ratio *R/K* is greater than or equal to a threshold (default: 0.05), i.e., this read contains a significant number of kmers and their reverse complement, we label it as potentially complex.

**Figure 2.**
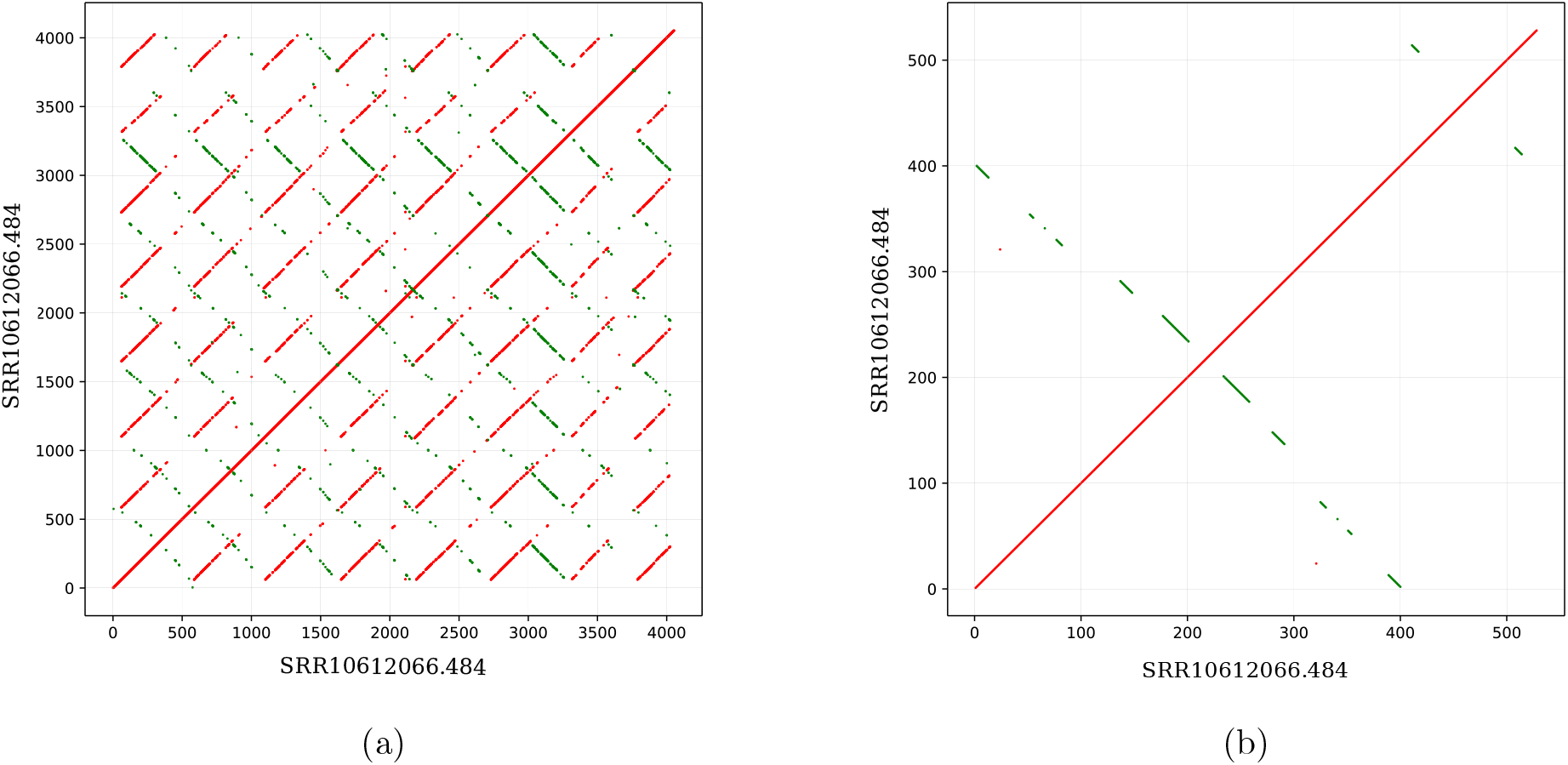
Dotplot of a complex read from the RCA protocol and its unit predicted by EquiRep. Red dots are shared kmers (length 10) between the sequence and itself; green dots are shared kmers (length 10) between the sequence and its reverse complement. (a) Dotplot of read SRR10612066.484; the dotted green lines show the occurrence of multiple repeating copies of the partial reverse complement of a potential circRNA sequence. (b) Dotplot of the unit predicted by EquiRep from SRR10612066.484.

### 2.2 Reconstruct repeat unit from potentially complex RCA long reads

Several existing tools can identify tandem repeats within a given sequence and reconstruct the sequence of the repeating unit. Some of these tools are specifically designed to tolerate sequencing errors. In our workflow, we use EquiRep to reconstruct the tandem repeat unit from each complex long read. Fig. 1b and Fig. 2b show the dotplots of the unit predicted by EquiRep from a regular and complex read respectively. The dotplot of the reconstructed repeat unit from a typical read is clean and shows no evidence of reverse complement signals. In contrast, the dotplot derived from the repeat unit of a complex read displays clear and prominent reverse complement patterns, providing preliminary support for our assumption. We denote the identified repeat unit from a potentially complex long read as *S*.

### 2.3 Conditional rearrangement of the repeat unit *S*

We hypothesize that the repeat unit *S* from a potentially complex long read contains a circRNA sequence followed by its partial reverse complement. Therefore, *S* can be represented as:

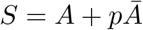

Here, *A* represents a nucleotide sequence, and *pĀ* denotes a partial reverse complement of *A* starting from the last base of *A*. To derive the algorithm, we divide *A* into 3 contiguous segments, *A* = *BCD*, where *B, C*, and *D* represent consecutive substrings of *A*. Correspondingly, *pĀ* consists of the partial reverse complement of the terminal substrings of *A*; we assume 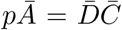. Thus, the full sequence *S* can be expressed as

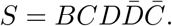

The predicted unit *S* can occur in five possible rotational configurations: 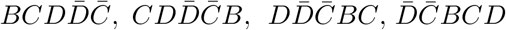, and 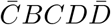, depending on the position at which the repeat unit begins (see Fig. 3). Regardless of the initial configuration, our objective is to rearrange *S* such that it begins with the partial reverse complement segment 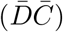, followed by the remaining portion of the repeat unit (*BCD*). To achieve this, we perform local alignment between *S* and its reverse complement 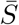. Based on the observed alignment patterns, the five possible configurations of *S* can be grouped into three distinct alignment categories.

**Figure 3.**
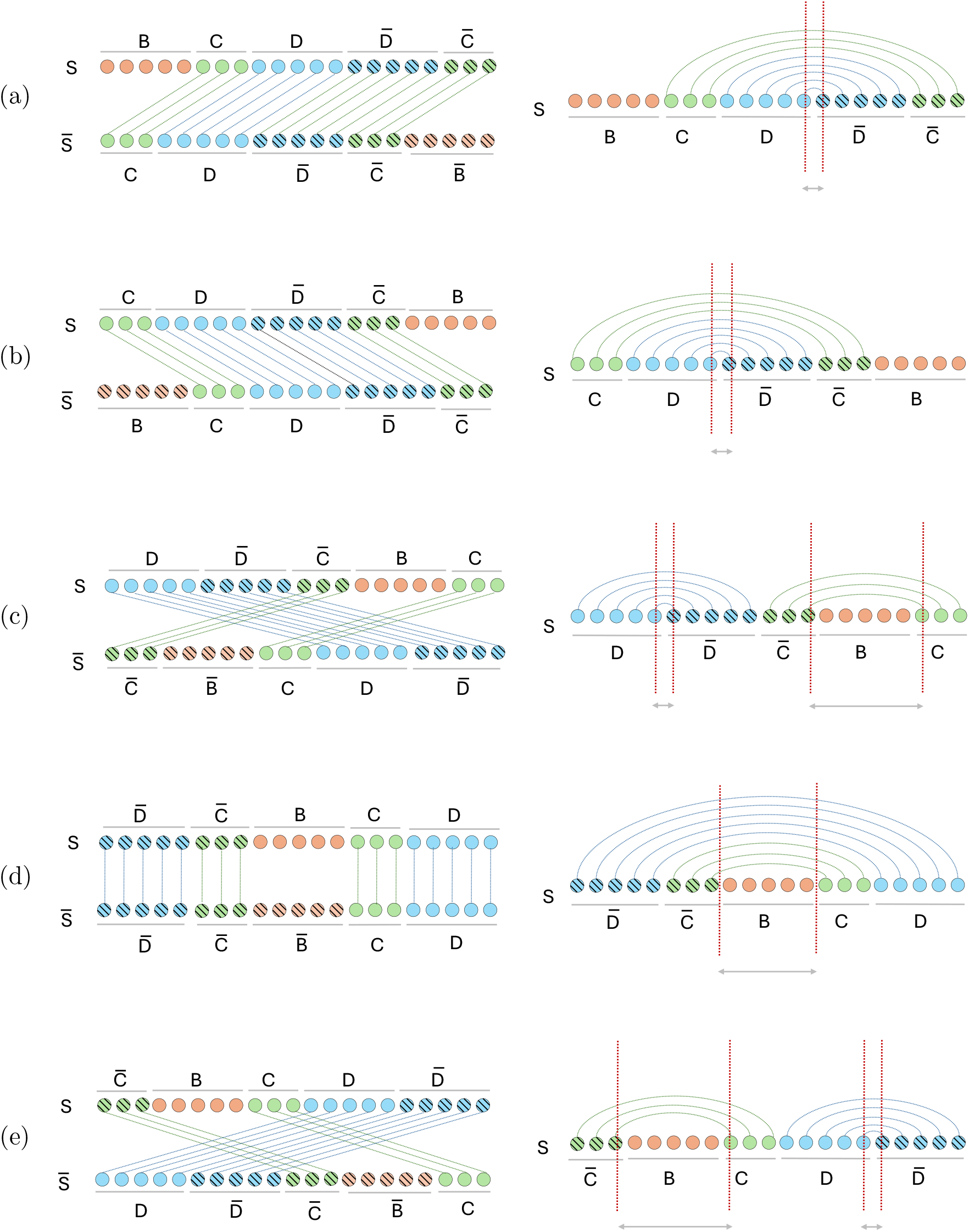
The predicted unit *S* can occur in five possible rotational configurations: (a) 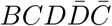, (b) 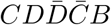, (c) 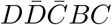, (d) 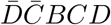, and (e) 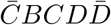. For each of the configurations, the left figure shows the local alignments between *S* and its reverse complement 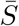; the right figure illustrates the aligned pairs arising from the local alignments (left figure). Each pair of red dotted lines represent the most inner pair (*x, y*) and the grey line marks the gap between *x* and *y*. In the cases where *S* needs rearrangement, the most inner pair with a very small gap (ideally 0) defines the boundary between 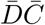 and *BCD*. For (d) where 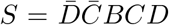, none of the most inner pair admits a small gap, suggesting that no rearrangement is needed.

In the first category, corresponding to the configurations 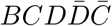 and 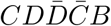, a single high-scoring local alignment is expected, where the segment 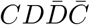 aligns to the same segment in 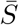 (Fig. 3a and Fig. 3b). The second category includes the configurations 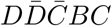 and 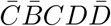, where multiple local alignments typically occur—specifically, between the segments 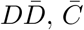, and *C* of *S* and their corresponding regions in 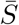 (Fig. 3c and Fig. 3e). In the third category, represented by the configuration 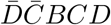, the sequence is already in the desired orientation, producing two local alignments: one between the segment 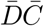 and its counterpart and another between segment *DC* and its counterpart in 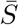 (Fig. 3d).

Assuming *n* is the length of *S*, an aligned pair between position *i* in *S* and position *j* in 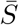 is equivalent to an aligned pair between position *i* and position *n* − *j* in *S*. For each alignment, we collect all aligned pairs in *S* and locate the pair (*x, y*), *x < y*, such that *y* − *x* is minimized; we call such pair as *the most inner pair* and define its *gap* as *y* − *x*. Observe that (see Fig. 3), for all configurations except 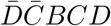 which is already the desired orientation, there is one local alignment where the gap between the most inner pair is very small, and that pair exactly separates 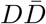 which is the right boundary to rearrange *S* to obtain the desired orientation.

Based on above hypothetical analysis, we design the following procedure to rearrange *S*. We perform local alignment between *S* and its reverse complement 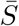. All resulting local alignment scores are recorded and ranked in descending order, and the top three non-overlapping alignment paths are selected. For each of the top three alignments/paths, we locate the most inner pair (*x, y*) and verify if *y* − *x < δ* (where *δ* = 2 by default). If such a pair is found in the highest-scoring path, the position *y* is treated as the rotation boundary, and *S* is rearranged such that it begins at position *y*. If no such pair is found in the top path, the second and third best-scoring paths are examined sequentially. If no such pair exists in any of the three paths, we conclude that the original sequence *S* is already in the correct rotational configuration.

### 2.4 Identify true circRNA sequence from the rearranged repeat unit

Assuming that *S* has been rearranged to the configuration 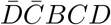 in the previous step, the next objective is to reconstruct the underlying circular sequence *A* from *S*, i.e., to separate the true circular sequence from its partial reverse complement. Two possibilities may exist: (1) the segment *BCD* represents the true circRNA sequence with 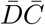 as its partial reverse complement, or (2) the segment 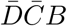 represents the true circRNA sequence with *CD* as its partial reverse complement. To identify both, we can perform global alignment between *S* and its reverse complement 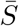. See Fig. 4. Since aligned pair (*i, j*) between *S* and 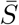 is essentially the aligned pair (*i, n* − *j*) within *S*, it suffices to only consider the region where *i < n* − *j*. In that region of the alignment table, the entry with the highest score must be from aligning 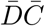 of *S* and 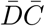 of 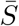, and that entry gives the exact positions to separate the true circular sequence and its partial reverse complement.

**Figure 4.**
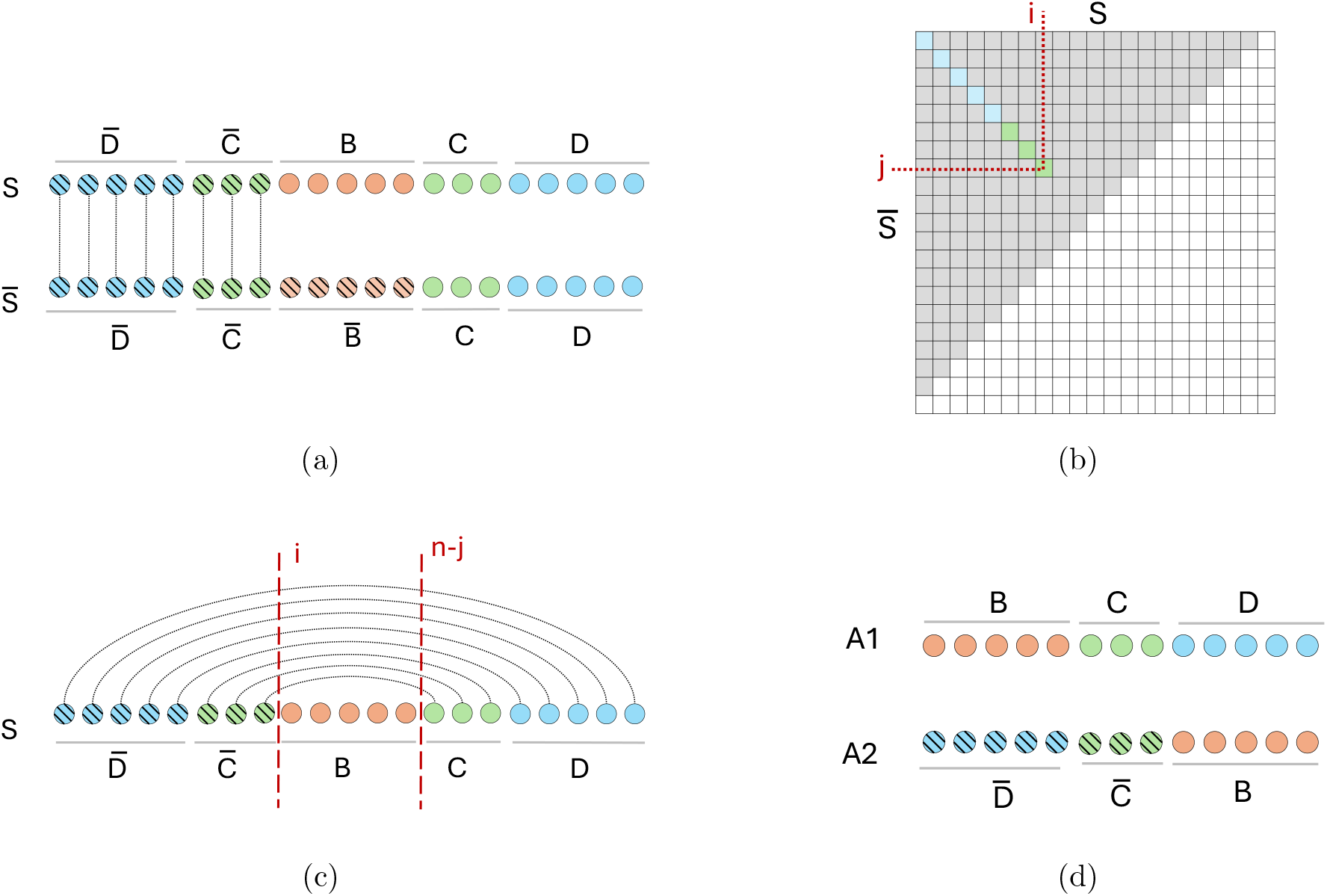
Identifying circRNA sequence from rearranged *S*. (a) Global alignment of *S* to its reverse complement 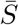. (b) We choose the highest-scoring entry (*i, j*) from the grey region *i* + *j < n* to identify partitions. (c) Determining partitions from the highest-scoring entry (*i, j*) in the global alignment, which are *i* and *n* − *j*, where *n* is the length of *S*. (d) The two possible circRNA sequences *A*_1_ and *A*_2_ derived from the partitions.

Based on above, we design the following procedure. We globally align *S* to its reverse complement. From the dynamic programming matrix, we identify the highest-scoring entry (*i, j*) that satisfies *i* + *j < n*, where *n* is the length of *S*. If *i/n <* 0.25 and *j/n <* 0.25, then the repeating unit *S* does not contain significant reverse complement and we do not take the originating read as complex. Otherwise, we label the read as complex read. In the latter case, we extract two substrings of *S, S*[1 · · · *n* − *j*] and *S*[*i* · · · *n*], denoted as *A*_1_ and *A*_2_. They are the two candidate circRNA sequences inferred from the input long read. There is no additional information in the long read to further infer which one is correct. CircPlex selects the final sequence based on alignment to a reference genome, described in Section 3.1.

## 3 Results

We evaluate the reconstructed circRNAs based on RCA long reads sequences of 2 human tissue samples (testis and brain; GEO accession number: GSE141693) from the isoCirc [39] protocol. The testis and brain samples comprise a total of 7,269,994 and 8,515,226 nanopore long reads respectively. We reconstruct putative circRNA sequences *A*_1_ and *A*_2_ for each of these reads, select one of them as the final circRNA based on some criterion, and evaluate our method in two ways: (1) comparison of circRNA sequence alignment to reference genome, (2) comparison of identified back splicing junctions to a circRNA database, circBase [14].

### 3.1 Comparison of circRNA sequence alignment to the reference genome

Alignment to reference genome has been used to assess the inferred circRNAs. Specifically, assume that the true circRNA sequence in the reference genome is *M*. Let *X* be a predicted sequence. Note that, *X* can be a rotation of *M* even if *X* is a correct prediction. To account for the rotation, one can concatenate two copies of *X* and aligns this concatemer to the reference genome. In the case that *X* is the correct circRNA (see Fig. 5b), *XX* must contain a substring, a rotation of *X*, that is the same (subject to mutations and sequencing errors) with *M*. This substring is therefore expected to be well-aligned to *M* and constitute the best/longest alignment between *XX* and the reference genome. In the case that *X* is incorrect, for example, *X* consists of the true circRNA and a partial reverse complement (see Fig. 5a), the best/longest alignment between *XX* and the reference genome is expected to be shorter than *X*. Denote the longest alignment as *X*_*M*_; we can use the 1 − |*X*_*M*_ |*/*|*X*| as a measure, termed over-prediction ratio, to evaluate the quality of *X*. An over-prediction ratio of 0 indicates a perfect prediction.

**Figure 5.**
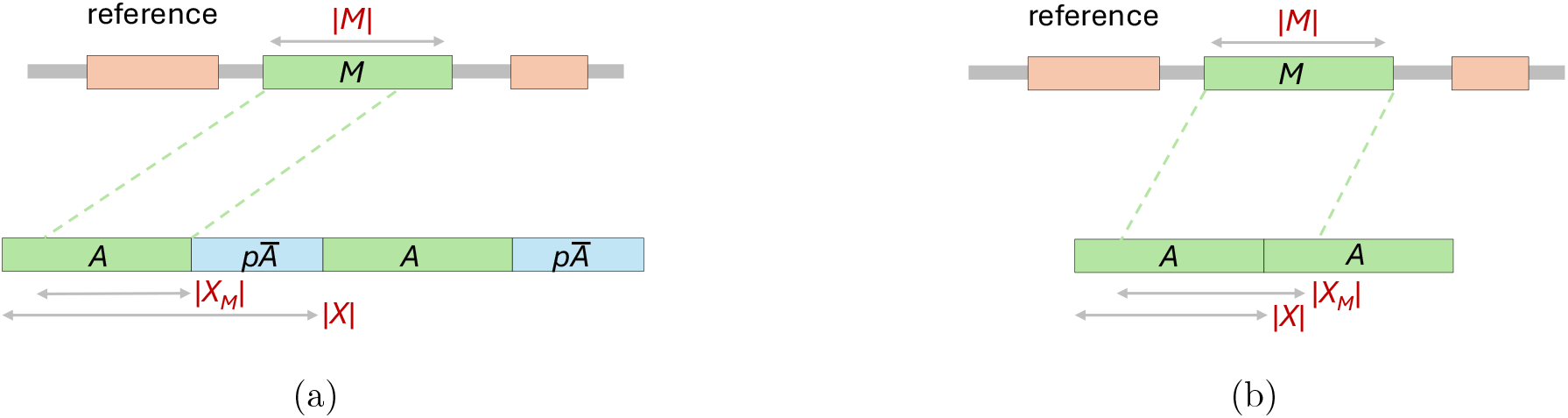
Evaluation of predicted circRNA sequence using alignment to the reference. The true circRNA sequence in the reference is labeled as *M*. (a) Alignment of concatenated copies of *S* (containing *A* and its partial reverse complement *pĀ*) to the reference. The unit length |*X*| is larger than the alignment length |*X*_*M*_ |. (b) Alignment of concatenated copies of circRNA seqeuence *A* predicted by CircPlex. The predicted circRNA length |*X*| is similar to the alignment length |*X*_*M*_ |.

Given a long read, let *S* be the repeating unit (i.e., the consensus sequence) predicted by an existing tool (in our case EquiRep is used). CircPlex predicts two candidate circRNA sequences, *A*_1_ and *A*_2_. All three predictions, *S, A*_1_, and *A*_2_, are assessed using above ratio. Specifically, we concatenate two copies of each sequence (*A*_1_ or *A*_2_) and align them individually. The concatemer is aligned to the reference genome using minimap2 [19]. The length of an alignment is computed by summing up the lengths of matching (M) and insertion (I) in the CIGAR string of an alignment record. To pick one from *A*_1_ and *A*_2_ as the final prediction, denoted as *A*, CircPlex chooses the one with longer alignment.

The distribution of the over-prediction ratio for *S* (i.e., the repeating unit) and for *A* (i.e., the CircPlex’s final prediction) over all complex reads in the testis sample is shown in Fig. 6. The plot for *S* shows a peak in the range 0.25-0.5, indicating that in majority of the long reads, the true circRNA sequences cover approximately 50%-75% of the repeat unit. In contrast, the distributions for CircPlex show more prominent peaks at zero, suggesting that a large portion our predictions correspond to correct, full circRNAs. Similar results are observed for the brain sample in Fig. 7.

**Figure 6.**
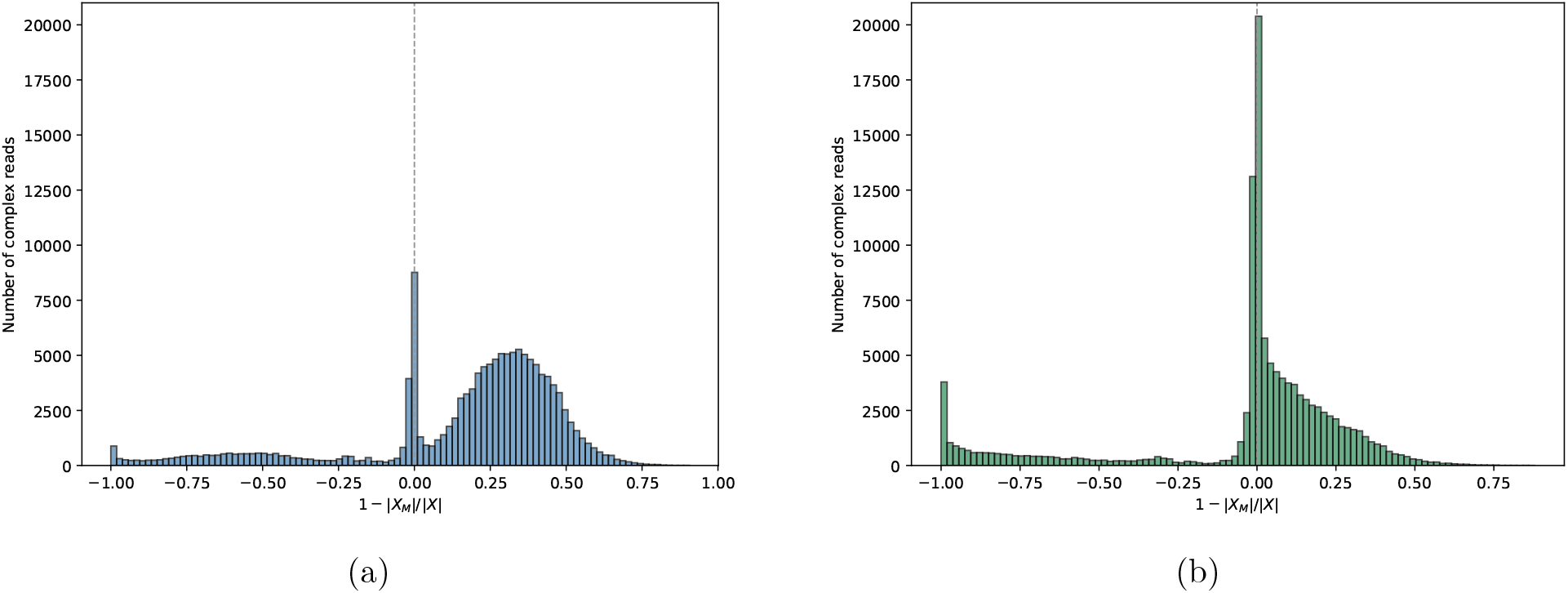
Distribution of over-prediction ratio from the testis sample. (a) Distribution for false prediction *S*. (b) Distribution for CircPlex prediction *A*.

**Figure 7.**
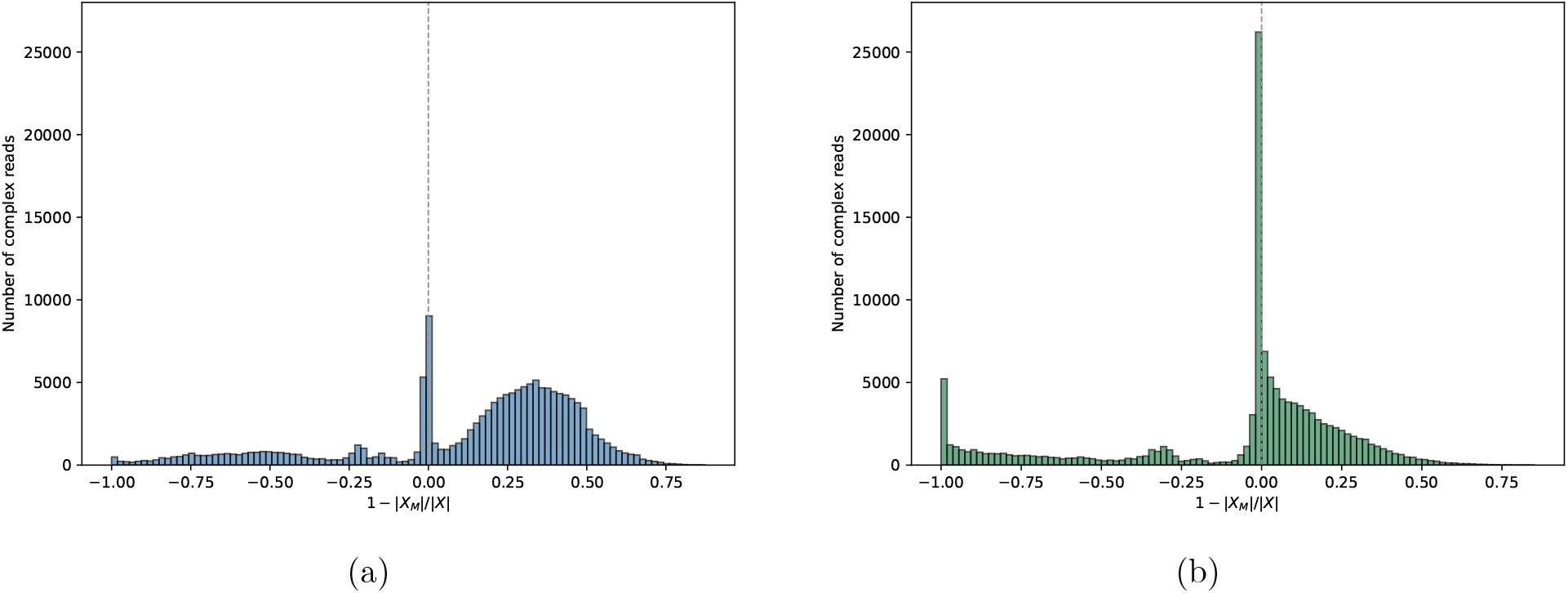
Distribution of over-prediction ratio from the brain sample. (a) Distribution for false prediction *S*. (b) Distribution for CircPlex prediction *A*.

### 3.2 Comparison of back splicing junction with database

Given a circRNA *A* predicted by CircPlex, we determine its BSJ from its alignment to the reference genome. Specifically, the BSJ coordinates of *A* are defined as the start and end positions of its alignment. Clustering is also performed: BSJs that originate from the same chromosome and both ends fall within 50 bp of each other are clustered together. Statistics on the CircPlex’s outcomes, including the number of complex reads, the number of BSJs and clusters, and the numbers restricted to complex reads with a over-prediction ratio in the range of [−0.1, 0.1], are reported in Table 1. We can observe that in both samples a substantial portion of reads are complex reads. Using minimap2 alignment, we detect 117,045 and 127,557 BSJs in the testis and brain samples, respectively, which are subsequently merged into 87,981 and 72,864 clusters. Notably, about 50% of the predicted BSJs and clusters correspond to circRNAs with absolute over-prediction ratio ≤ 0.1, supporting our hypothesis and validating the derived circRNA sequences by CircPlex.

**Table 1:** Summary of CircPlex outcomes on both samples.

| Count | Testis | Brain |
| --- | --- | --- |
| # Reads | 7,269,994 | 8,515,226 |
| # Complex Reads | 784,365 | 1,394,188 |
| # BSJs | 117,045 | 127,557 |
| # BSJs with over-prediction ratio in $[-0.1, 0.1]$ | 58,334 | 57,441 |
| # Clusters | 87,981 | 72,864 |
| # Clusters with over-prediction ratio in $[-0.1, 0.1]$ | 46,340 | 38,435 |

We report the BSJs derived from the circRNAs predicted by CircPlex and evaluate their overlap with isoCirc and circBase at both the read level and the cluster level. We also evaluate overlaps for the subset of CircPlex circRNAs whose absolute over-prediction ratio is less than 0.1, i.e. circRNAs that are strongly supported based on the evaluation described in Section 3.1. Two BSJs are considered overlapping if they occur on the same chromosome and their positions are within 50 bp of each other. The results of overlaps for the testis and brain sample are illustrated in Fig. 8 and Fig. 9 respectively. We observe that in all case, CircPlex identifies a significant number of circRNAs that overlap with circBase but are unidentified by isoCirc. About half of these circRNAs have absolute over-prediction ratio ≤ 0.1, providing strong evidence that the circRNAs we identified are genuine.

**Figure 8.**
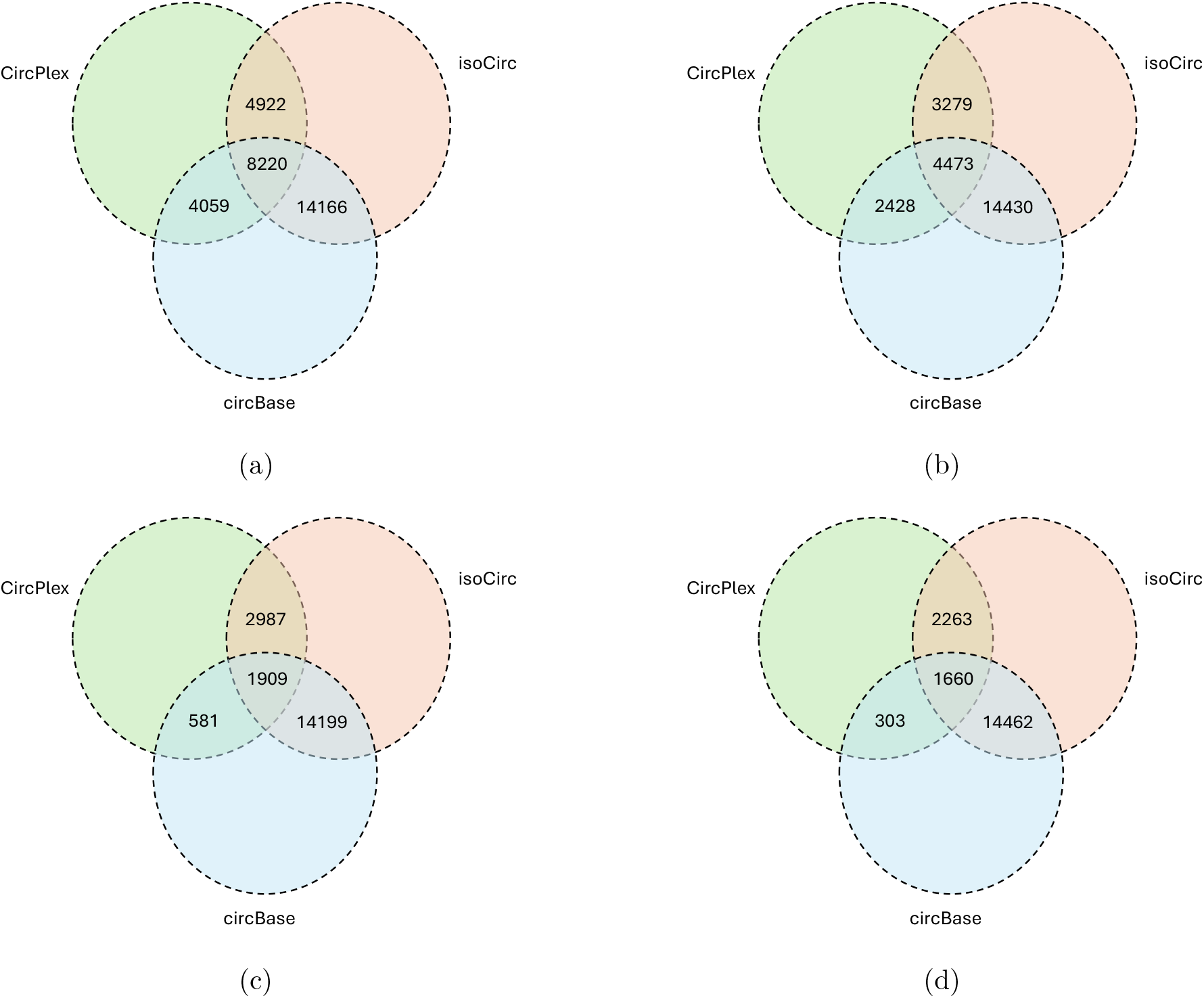
Overlaps of BSJs from CircPlex and isoCirc with circBase for the testis sample. (a) Overlap at read level for circRNAs predicted by CircPlex. (b) Overlap at read level for circRNAs predicted by CircPlex with absolute over-prediction ratio ≤ 0.1. (c) Overlap at cluster level for circRNAs predicted by CircPlex. (d) Overlap at cluster level for circRNAs predicted by CircPlex with absolute over-prediction ratio ≤ 0.1.

**Figure 9.**
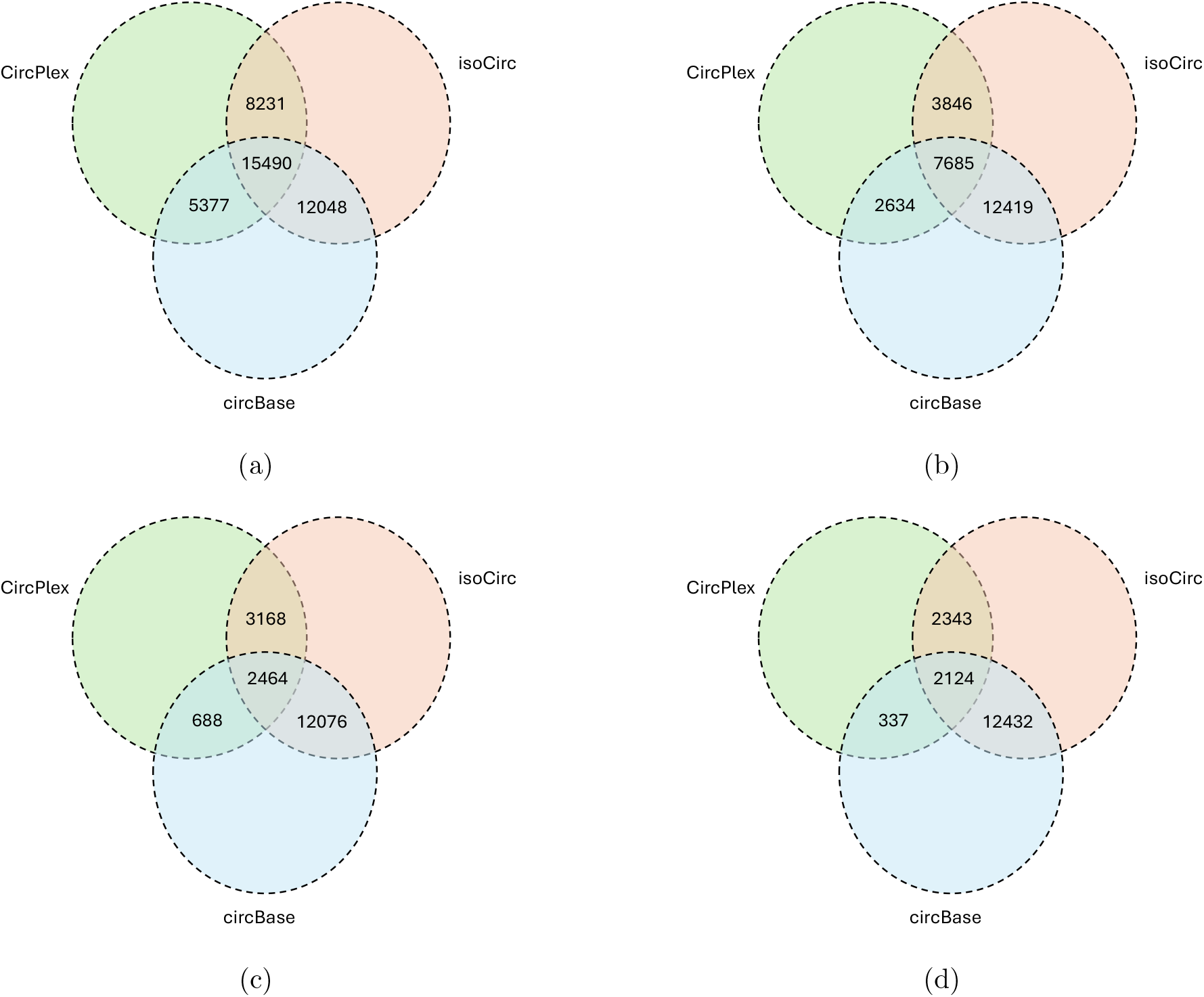
Overlaps of BSJs from CircPlex and isoCirc with circBase for the brain sample. (a) Overlap at read level for circRNAs predicted by CircPlex. (b) Overlap at read level for circRNAs predicted by CircPlex with absolute over-prediction ratio ≤ 0.1. (c) Overlap at cluster level for circRNAs predicted by CircPlex. (d) Overlap at cluster level for circRNAs predicted by CircPlex with absolute over-prediction ratio ≤ 0.1.

## 4 Conclusion and Discussion

Circular RNAs have emerged as a topic of significant interest in the scientific community due to their unique properties and diverse potential applications. The rapidly expanding research in circRNA biology relies on accurately determining their full-length sequences. However, precise identification of complete circRNA sequences remains a challenging problem. In this study, we present CircPlex, a method designed to assemble circRNAs from complex rolling-circle sequencing data. Our analysis reveals that the repeating units within the long reads sequenced from the RCA protocol often contain not only the circRNA sequence but also its partial reverse complement. Building on this observation, we introduce an approach to extract the true circRNA sequence from the full repeat unit. CircPlex takes a long read as input and uses a kmer-based approach to first identify potentially complex reads. It then applies local alignment on the repeating unit to rotate it into a specific orientation. Finally, a modified global alignment approach is used to extract the circRNA sequence embedded within the unit. The resulting circRNA predictions are validated using alignment and comparison of BSJs to existing database. Results strongly support our hypothesis that repeat units contain partial circRNA sequences and confirm the reliability of the circRNAs extracted by CircPlex, many of which were missed in the isoCirc pipeline.

While our findings are promising, we recognize that errors in upstream processing or limitations of the EquiRep pipeline can propagate into downstream computations. Future extensions of this work could focus on designing a tool that directly identifies circRNA sequences from multiple copies in long-read data, bypassing the need for intermediate repeat detection steps. Although we generated a large number of circRNA sequences, not all may represent true circRNAs. A machine learning model could be developed in future to distinguish true circRNAs from false positives using sequence patterns derived from our predictions. We observed relatively low overlaps between our identified BSJs and circBase annotations. This discrepancy is likely due to the incompleteness of circBase and suggests that many circRNAs remain largely unannotated. Consequently, the circRNAs identified by our method may help refine existing databases and expand the catalog of known circRNAs. Overall, CircPlex demonstrates strong potential for discovering novel circRNAs and provides a useful framework for full-length circRNA assembly. We are optimistic that further refinements and complementary approaches will enhance applicability of this method in circRNA research.

## Availability

The source code of CircPlex is freely available at https://github.com/Shao-Group/CircPlex. The scripts, evaluation pipelines, and instructions that can be followed to reproduce the experimental results of this work is available at https://github.com/Shao-Group/CircPlex-test.

## Acknowledgment

This work is supported by the US National Science Foundation (2145171 to M.S.) and by the US National Institutes of Health (R01HG011065 to M.S.). We thank Xiaofei Carl Zang for helpful input.

